# PAM-DB: Revealing Protein Activation Mechanisms for Next-Generation Rational Drug Discovery

**DOI:** 10.64898/2026.08.20.745895

**Authors:** Xiaohong Zhu, Xiangyu Li, Yaning Hou, Ruifeng Zhou, Yingchao Yan, Arieh Warshel, Chen Bai

## Abstract

Current rational drug design relies predominantly on computational (CADD/AIDD) methods that model binding thermodynamics and static conformations of target proteins, primarily in their inactive states. However, the kinetic parameters that govern experimental efficacy—such as catalytic turnover and signaling potency—are determined by molecular interactions with transition states (TS), intermediate states (IS), and the entire continuum of conformations along the least free-energy activation pathway. The absence of this dynamic dimension has fundamentally limited the predictive power and success rate of conventional structure-based approaches. Here, we present a structural database that systematically maps the complete activation trajectories of pharmaceutically relevant targets, encompassing TS, IS, and all connecting conformational ensembles. This resource offers multiple strategic advantages for drug discovery: enabling rational targeting of previously “undruggable” proteins, facilitating biased agonism/antagonism design, revealing cryptic allosteric sites in inactive conformations, identifying novel transient pockets along the activation route, rationalizing the mechanisms of existing drugs, predicting mutational effects on activation barriers, and prospectively forecasting drug resistance and off-target liabilities. We demonstrate the utility of this database through representative case studies and provide implementation guidelines for integration into existing discovery pipelines. More detailed information can be found at our website: https://www.momedpamdb.com/en.

**Terminology:** The following terms are clarified in this document:

Stable state (SS): In this document, this term refers exclusively to, and is synonymous with, the protein’s inactive state (IAS). Note that other states may also be stabilized into meta-stable states by certain means.

Unstable state (US): This term encompasses all states other than SS, even if they appear computationally meta-stable on the free energy surface.

Activated state (AS): The meta-stable working state of the protein.

Transition state (TS): The state with the highest free energy along the least-energy pathway on the free energy surface that connects the inactive state to the activated state of the target protein.

Intermediate state (IS): The state(s) located at a local minimum along the least-energy pathway, excluding SS and AS.

## Introduction

Computational and AI-based tools for studying protein structure, energetics, and molecular binding patterns have become indispensable in modern rational drug design. Molecular docking, sampling algorithms (molecular dynamics, Monte Carlo, etc.), potential functions (quantum mechanics, molecular mechanics, etc.), free energy calculations (FEP, TI, etc), and virtual screening are being used in almost all relevant organizations prior to committing to wet-lab experiments. These computational data, together with experimental results are also being used by AIDD industry to fuel AI generative or predictive models.

However, such efforts are almost exclusively focused on the inactive state (IAS), and in some cases also the active state (AS) when experimental structures are available (AI models such as AlphaFold or RoseTTAFold are sometimes used to generate stable-state structures as well). Practically, this remains the only efficient way to provide structure–activity relationship (SAR) information to computational biologists, medicinal chemists, and AI models—even though it is not necessarily scientifically complete.

In most cases, functional proteins reside predominantly in the stable inactive state (IAS), but it is the transition process from the IAS to the AS that determines their kinetic parameters and macroscopic functions. As shown in the simplified activation coordinate diagram in Figure 1, when the target protein transitions from the IAS to the AS, it undergoes a process in which the free energy first rises and then falls, accompanied by substantial conformational changes, during which the system may pass through multiple kinetically significant states. For simplicity, only a single transition state (TS) is shown in the figure. Along this path, the point with the highest energy is defined as the transition state (TS), and the energy difference between the TS and the IAS defines the activation barrier. Additional intermediate states (ISs) or other TSs may also exist along the pathway, but they do not affect the definition of the overall barrier or the discussion here. When a drug molecule binds to the target protein, if it raises this barrier, it inhibits the transition from IAS to AS and thereby impairs protein function, thus serving as an inhibitor; conversely, if it lowers the barrier, it facilitates the activation process and acts as an agonist. Therefore, understanding the complete activation process of the target protein and how candidate drugs affect this process is essential.

**Figure 1.**
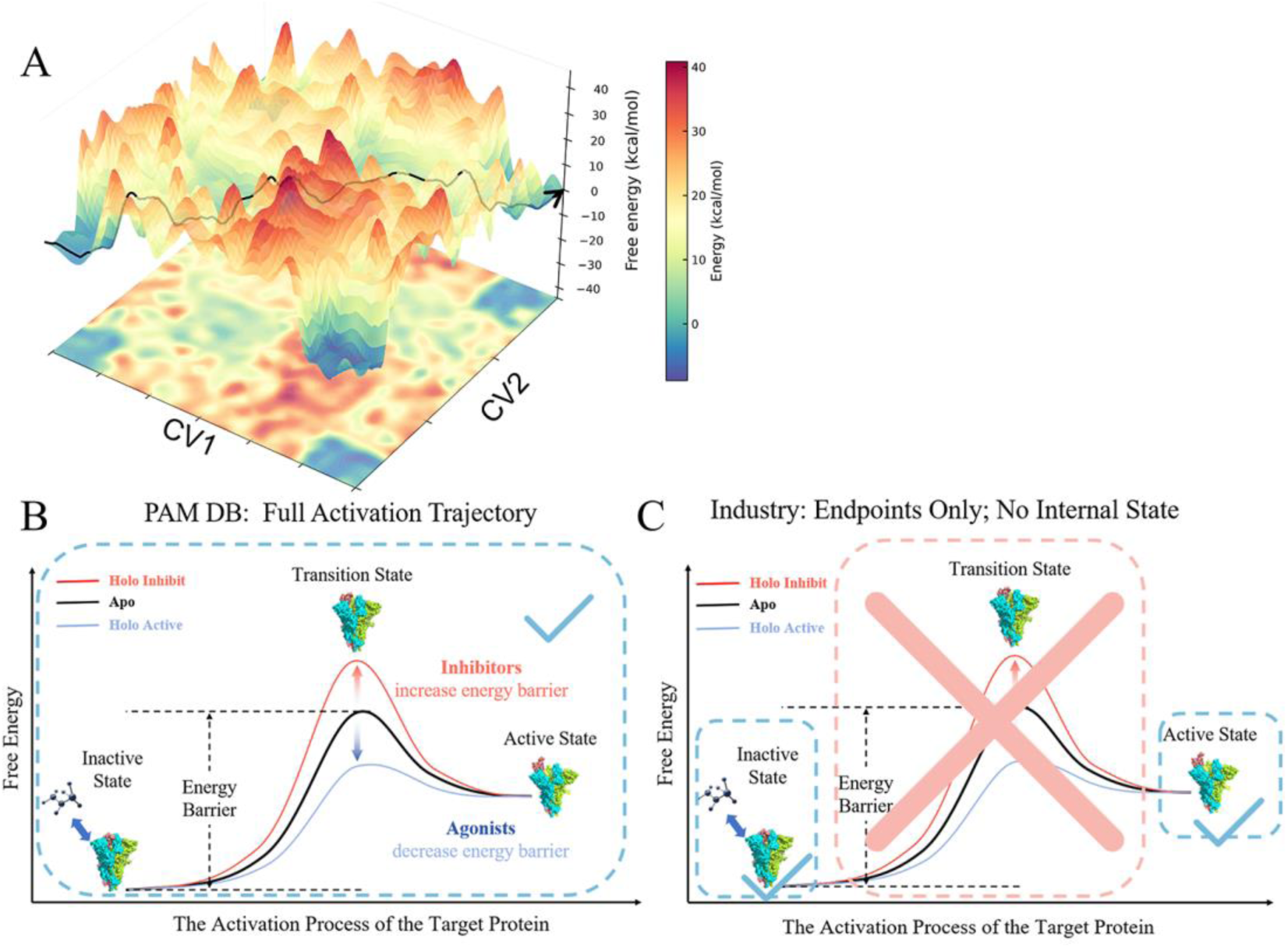
(A) The Example of a multidimensional protein activation free-energy landscape is projected onto a representative one-dimensional activation coordinate, from which the simplified curves in panels B and C are derived with least free energy pathway depicted in black line. (B) The black line indicates a simplified activation coordinate of a target protein, with IS and other TS not shown. The red line and blue line show how inhibitor and agonist molecules affect the activation barrier. Our database captures the complete activation process of the target protein, including intermediate states (IS), TS, the states connect them, and their associated kinetic information. (C) Industry studies are generally limited to the endpoint stable structures (IAS and some AS), leaving the intervening conversion pathway inaccessible.

The current database provides the complete conformational changes and energetic information for a set of target proteins throughout their entire activation process. This kinetic perspective is not merely a theoretical construct. It is grounded in our decade-long systematic benchmarking of computed barrier heights and activation landscapes against experimental rate constants across multiple protein families. This extensive calibration ensures that the activation pathways presented in this work are not speculative trajectories but physically faithful representations that withstand experimental scrutiny. In contrast, traditional CADD or AIDD workflows are limited to studying only the endpoint structures, whereas the internal conformational changes, energetic profiles, and other key kinetic details remain a black box.

Without activation information, drug design is akin to crafting a key for a lock without knowing the lock’s internal mechanism (Figure 1C). One could design many keys that fit the keyhole, but whether they can actually open the lock after insertion is largely a matter of chance—since the internal structure of the lock remains unknown. Traditional docking techniques operate under the same logic: one can design or screen a vast number of molecules that bind strongly to the IAS, but their effects on the internal conformational transitions—especially on the TS and on the energy barrier—remain entirely unknown. This, we believe, is a major reason why the success rate of conventional CADD and AIDD remains relatively low.

Due to experimental limitations, state-of-the-art techniques such as cryo-EM or X-ray crystallography cannot directly capture the internal structures that connect the SS and the AS. Only in some cases can the AS be solved by adding stabilizing agents (e.g., ligands or nanobodies), thereby trapping the complex in a metastable conformation. Recent large-scale HDX-MS studies^1^ have begun to profile conformational fluctuations of small proteins domains of tens of amino acids, yet the direct structural characterization of intermediate and transition states remains scarce. At present, the only practical way to obtain intermediate states, transition states, and the conformations between them is through non-equilibrium computational modeling.

However, from computational perspective, many obstacles remain. First of all, ASs have high free energy and low occurrence rate. Classical sampling methods such as ordinary molecular dynamics (MD) simulation and Metropolis Monte Carlo (MC) algorithm will eventually converge to the canonical ensemble distribution, which clusters around the energy basin bottom. Thus, these methods are not suitable for capturing high-energy states. Enhanced sampling methods do exist-such as TMD, replica exchange MD, metadynamics, etc. Yet they are still difficult to apply in practice. A typical biophysical system contains thousands of amino acids (hundreds of thousands of atoms, corresponding to 3N degrees of freedom). Even a small translational displacement of a single helix can cause severe atomic collisions and simulation failure, let alone large-scale domain motions or rotational rearrangements.

For example, in previous work on β2AR^2^ (Figure 2), we found that upon agonist binding, the receptor opens a cavity on the cytoplasmic side, allowing the Gs α5 helix to intrude, straighten, and rotate. Subsequently, Gs protein undergoes a large-scale rearrangement in which the Ras-like domain separates from the α-helical domain, opening the nucleotide binding pocket. These motions involve coordinated displacements across multiple domains over distances of tens of angstroms—far beyond the scale of local fluctuations that conventional sampling methods can reliably capture. Similarly, for ATPases^3^, the top α₃β₃ ring undergoes sequential opening and closing motions coupled with stepwise rotation of the central stalk, with each ∼120° rotation requiring concerted rearrangements across multiple subunits. The sheer scale and coupling of these conformational changes illustrate why enhanced sampling methods alone are often insufficient and why specialized workflows are needed.

**Figure 2.**
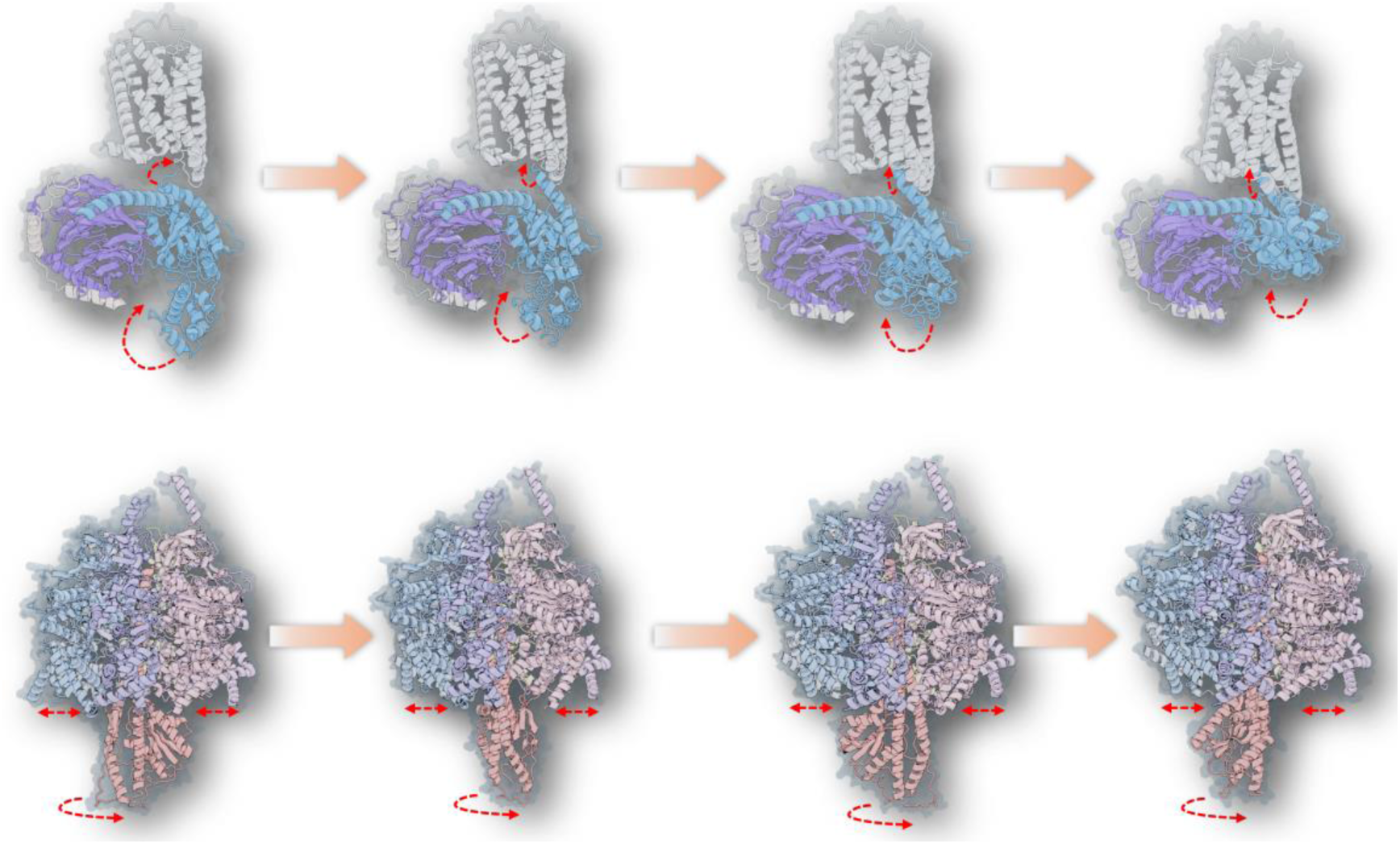
Conformational changes during the activation process of β2AR (top panel) and chloroplast F1-ATPase (bottom panel).

Second, the transition from the SS to the AS is usually not a linear process; one cannot simply “drag” a protein or complex by applying an external force or minimizing the structural RMSD between the IAS and the AS. In multimeric complexes, the coupling and relative movement between different monomers are challenging to model, as the conformational change of one monomer can be sterically constrained by another.

For example, in our previous work on the class C GPCR metabotropic glutamate receptor 2 (mGlu2) homodimer^4^, cryo-EM had resolved the endpoint states—yet molecular and energetic details of the mGlus activation remain elusive. A Targeted MD attempt to directly connect IAS and AS led to steric collisions between the two 7TM domains of the dimer (Figure 3A), revealing that the rotational motions of the two subunits are coupled rather than synchronous, a feature that simple “drag” approaches cannot handle.

**Figure 3.**
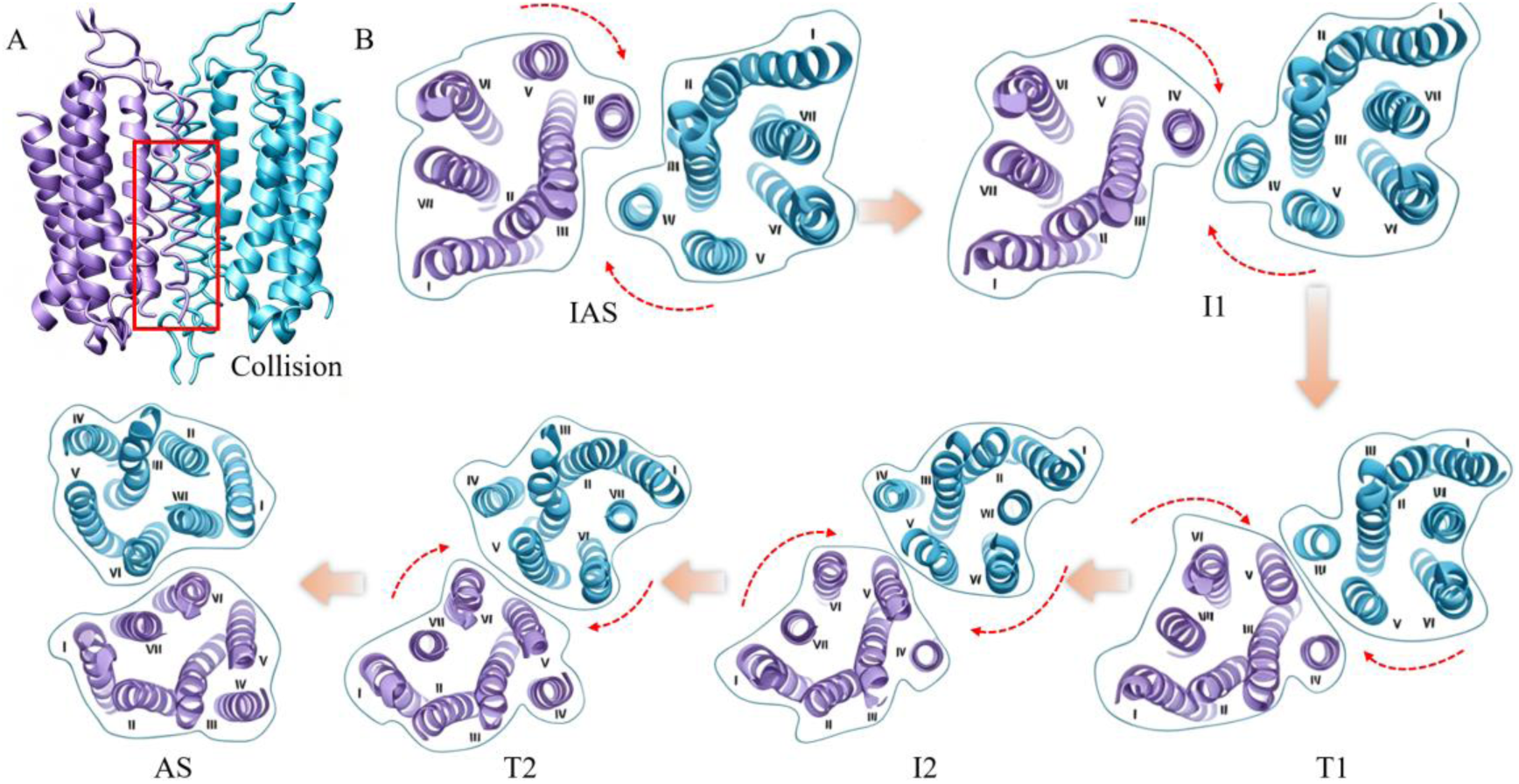
(A) Steric collisions occur between the 7TM domains during TMD. (B) The complete conformational transition from IAS to AS captured by our method, demonstrating that the rotational motions of the two subunits are coupled rather than synchronous. Multiple dimerization structures of the mGlu2 transmembrane domain (the extracellular view) from the IAS to AS state, with the red arrows indicating the direction of 7TM rotation. The ATPase example in Figure 2 holds the same logic.

Third, coupling between different types of events—such as conformational change, ligand binding/release, and chemical bond formation/breaking—poses additional challenges. We have previously tackled such multi-event coupling in many specific systems.^2, 4–8^ For example, in the GPCR family, we studied the coupling between receptor conformational changes, G protein binding, and GDP release/dissociation.^2, 4, 5^ We also investigated spike-host receptor binding coupled to conformational rearrangement in SARS-CoV-2.^8^ Other examples include conformational changes coupled to ion transport in ion channels.^6, 7^ All these cases involve the coordination of multiple events—conformational changes, ligand binding/release, and chemical transformations—whose interplay can only be fully understood through a multi-dimensional free energy landscape and the associated minimum free energy pathway.

All these issues must be addressed before a reasonable multi-dimensional free-energy landscape can be obtained. From this landscape, the minimum free-energy pathway can then be extracted, which is essential for understanding the activation mechanism of biophysical systems and, subsequently, for evaluating how drug candidates affect it.

The third difficulty mentioned above is important in academic studies but is not indispensable for ordinary CADD or AIDD use of current database. See the instructions below.

## Discussion

Studying the activation process of target proteins is already a challenging topic in academia, and the difficulty level increases substantially when we attempt to generate a standardized dataset that can be directly used by ordinary practitioners in the industry, while maintaining reasonable quality control and ease of use. We have worked in this field for years and have developed a reliable and robust procedure to generate the full protein/complex coordinates of the structures along the transition between the SS (IAS) and the AS. ^2–12^ The current work should fulfill the needs of most users.

For validation, even though TS structures cannot be directly captured experimentally, we can still indirectly validate the modeling results by comparing calculated energy barriers with experimentally measured reaction rate constants using kinetic relationships such as the Arrhenius equation. See our previous works.^2–12^ We also show several examples later in this document of how we utilized this dataset to identify novel pockets, design new molecules, and optimize existing ones.

This is not a mathematically exact solution in the way that quantum mechanics applied to few-particle systems would be, but as mentioned above, it arguably represents the most reliable and reasonable approximation of the activation process of functional proteins at the current stage. It should be able to provide valuable information for understanding how proteins work, for enhancing CADD procedures, and for generating training data for AIDD models.

Here we present the first version of PAM-DB v1.0 (Protein Activation Mechanism Database), which provides All Atom-level coordinate details of the entire activation process for multiple important target proteins. We will continue to update the database and include more proteins over time. Although different computational groups have different levels of modeling expertise and may adopt various modeling strategies, here we provide a “standard” dataset that can be used by a typical computational biologist or pharmaceutical chemist from any institution or company. At the same time, we encourage users to leverage their own expertise to help the database and workflow evolve.

We offer the following suggestions on how to utilize this dataset to facilitate molecular design.

### Usage instructions

1. Track how the binding pocket reshapes throughout the activation process. Since protein structures undergo substantial conformational rearrangements during activation, this analysis can reveal previously uncharacterized binding sites and reshaping of old pockets— information that is particularly valuable for targeting undruggable proteins or designing allosteric drugs (Figure 4). This is in contrast to long MD simulations near the IAS. The new pocket information could be directly used in CADD procedures, such as screening.
2. Understand how old molecules work before designing new ones when performing optimization. Identify binding features—such as binding pose, key interactions, binding free energy—of ligands with Uss through activation process, even though they might already exhibit strong binding affinity in the SS. This procedure is very useful in designing biased agonists/antagonists or optimizing existing molecules by manipulating key interactions. We would like to emphasize here one should not rely solely on the binding pose and binding energy of molecule at internal states that are obtained from docking software. A more meaningful and reasonable practice is to find binding poses of molecule of each internal state that can lead to a consecutive movie.
3. Calculate the ligand effect on energy barrier ΔΔ*G*, instead of only focusing on binding energy at SS (IAS) during molecule evaluation. An economic way is to obtain this value by examining only the IAS and the TS, that is, calculating the new energy barrier of the holo process after adding the binding energy difference between the TS and the IAS. Then deduct it from the conformational barrier value of the apo process (Eq. 3), see example 2 (<u>Biased antagonist design for an undruggable target</u>). Otherwise, the computational cost would be massive since binding effect of every holo state needs to be re-calculated through the activation pathway, in order to get a more accurate drug effect on barrier height. Fortunately, the conformational energy contribution could be canceled out in measuring ΔΔ*G* and computational biologist could focus on binding calculation (Eq.6). Let *G_IAS_*_,*apo*_, *G_TS_*_,*apo*_, *G_IAS_*_,ℎ*olo*_, and *G_TS_*_,ℎ*olo*_be the free energies of the four states, and let *G_L_* be the free energy of the free ligand. In reality, the TS conformation differs between the apo and holo states; however, as a simplifying approximation, we assume that the TS state position does not change during a holo process. The activation barrier without ligand is

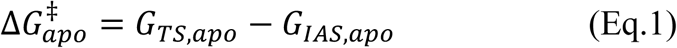

and with ligand it is :

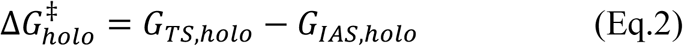 Therefore, the effect of the ligand on the entire activation process (the change in activation barrier) is

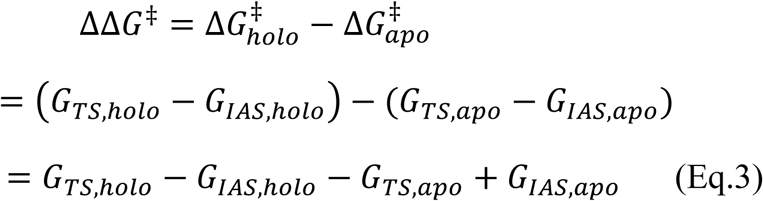

On the other hand, the binding free energies of the ligand to the IAS and TS are:

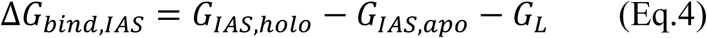

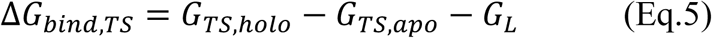

Thus, the difference in binding free energies between the two states is:

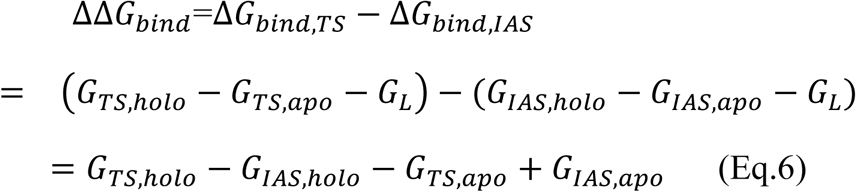

Comparing Eq.3 and Eq. 6, their expressions are identical, therefore:

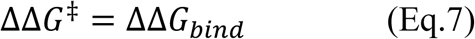

This result shows that the effect of the ligand on the activation barrier is numerically equivalent to the difference in its binding free energies to the TS and IAS. Since the free energy of the free ligand *G_L_* cancels out completely during subtraction, we do not need to compute the absolute solvation free energy or conformational entropy of the ligand in solution. Consequently, the entire problem reduces to calculating the binding energy difference ΔΔ*G_bind_*, which is precisely why computational biologists can focus on binding calculations rather than on the complex conformational energetics of the protein. When searching for agonists, one should not rely on a single metric, for example, simply checking whether a molecule binds stably to the AS. In fact, there are two common mechanisms for agonist action. In the first mechanism, the agonist should have decent binding affinity to both the IAS and the TS, while simultaneously lowering the activation energy barrier. In the second mechanism, the agonist should bind strongly to the AS and raise the backward energy barrier, thereby stabilizing the active state and prolonging its residence time. For antagonist design, a steric antagonist should bind strongly to the IAS while blocking the activation process through steric collision. For a non-steric inhibitor, the binding energy between the antagonist and the TS or AS should be more positive than that between the antagonist and the IAS, thereby increasing the overall activation energy barrier or the reaction energy. This third usage is useful in designing and evaluating novel skeletons but is a little bit tricky; for non-specialists, we suggest just using the first two strategies.
4. Use your own molecule database (small molecule, peptide, or any molecule type that might fit the binding pocket) to perform CADD screening by incorporating the information and criteria obtained from steps 1–3 or one of them.
5. Construct and train new AIDD models or adjusted old AIDD models by incorporating information from steps 1-4 or one of them.
6. Select molecule candidates that exhibit favorable features identified in steps 1–4 or one of them for further experimental validation, in addition to your traditional CADD/AIDD criteria.
7. Using the information obtained from steps 1–4, one can also compute the effect of known ligands on the activation energy barrier (ΔΔG) for a given target. These computed ΔΔG values can then serve as a benchmark to evaluate old molecules, generate positive and negative data to refine existing AI models.
8. Other than ligand binding effect, the current database can also predict drug resistance and off-target effects. It enables the simulation of how mutaionals affect protein activation, allowing for prospective prediction of drug resistance. Simultaneously, by modeling drug binding to different conformational states, it facilitates the assessment of potential off-target effects, thereby enhancing safety profiles.

**Figure 4.**
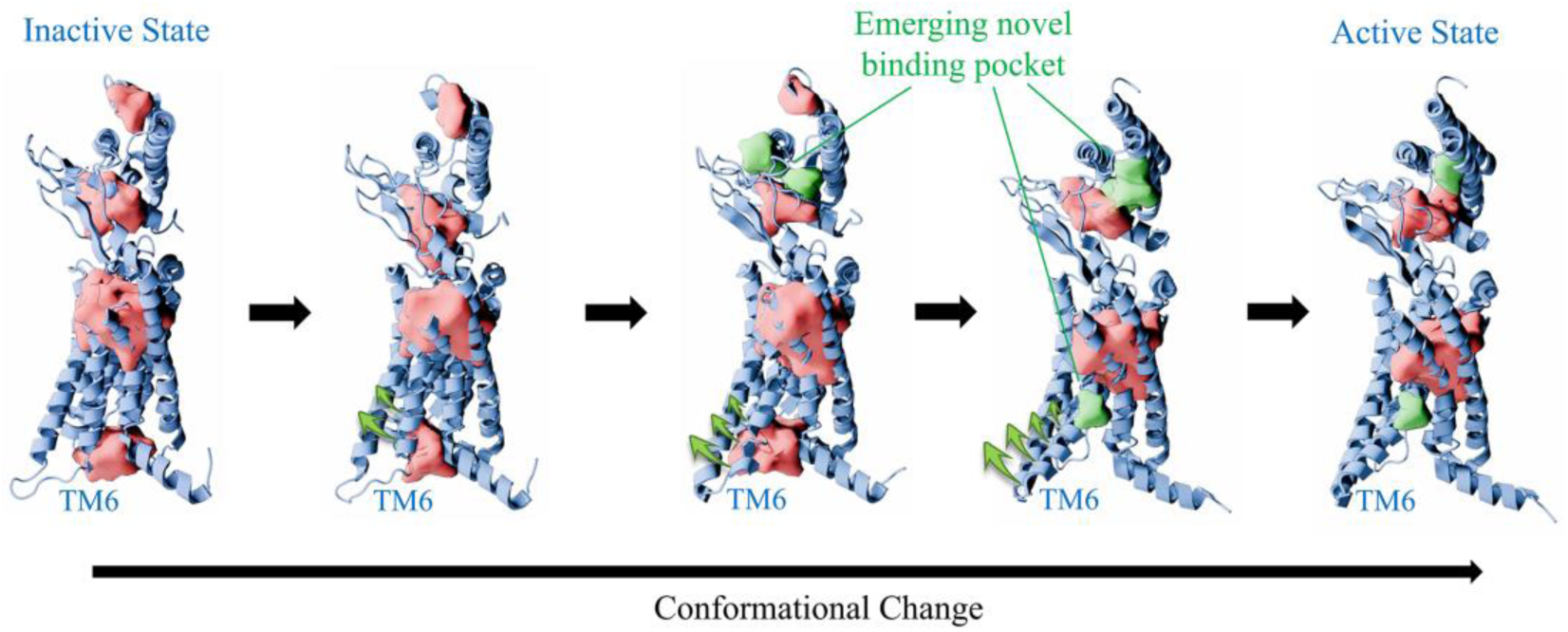
Conformational transition of AMY3R and the selective emergence of a novel pocket. This diagram illustrates the structural trajectory of the AMY3R receptor transitioning from its inactive state to the active state. Throughout the activation process, the orthosteric pocket in the transmembrane domain (lower red volume) remains relatively stable, exhibiting only minimal local variations. Concurrently, the non-transmembrane extracellular domain (ECD) pocket (upper red volume) undergoes a dynamic spatial relocation in response to the unfolding and expansion of the extracellular helices. On the intracellular side, receptor activation is driven by the pronounced outward rotation and displacement of transmembrane helix 6 (TM6, as indicated by the green arrows). This dramatic helical movement facilitates the gradual opening and exposure of a transient, state-specific “novel pocket” (depicted in green) near the cytoplasmic face.

The above suggestions represent straightforward applications of the current dataset. However, users are encouraged to perform customized analyses according to their own research needs and expertise.

### Precautions for dataset use

The unique feature of this dataset is that it is not a classical sampling trajectory such as ordinary trajectories, in which unreasonable bond distances are relaxed and corrected with a proper set up. However, to get the consecutive coordinates that connect the SS-(IS-TS)n-AS process, we need to go uphill in energy surface for a large amount of residues. Such an uphill process inevitably renders some interatomic distances unreasonable from the perspective of a classical modeling. This could decrease the readability of the data for computational software to a large extent. In practice, when using our raw untrimmed structures (as in the basic version) for subsequent simulations, the program will show many error messages due to too long or too short interatomic distances, making it almost impossible to model or to compute energies.

For non-expert users, we have simplified the modelling process by providing trimmed structures (in the standard version) that could be directly used in docking and modelling. One needs to keep in mind, for the AS, we need to constrain/freeze the degrees of freedoms (DOFs) of high energy area to account for the environment effect, while keeping the binding pocket flexible.

For experienced users who want to play with the raw data, we give the following suggestions according to our experiences:

1. To obtain the conformational change free energy of a target protein through the activation, we suggest converting the all-atom coordinates into a CG representation (Figure 5) while preserving the backbone spatial distribution, then sampling with the main-chain freezed or relaxed with restraint force, otherwise the conformation might exit the high-energy state. In previous academic works, we utilized a non-free package Molaris-XG. The Molaris force field could yield reasonable activation barrier height that is comparable to experimental rate constant value. However, in this dataset, we would like to switch to an open source force field Martini. We performed benchmark between Molaris and Martini force field and found Martini systematically amplify the barrier heights, in contrast to Molaris. For a more rigorous conformational free energy calculation, one could perform FEP calculation on a lot more internal states with good phase space overlap. However, such expensive calculation is out of interest here since the conformational contribution is canceled out. (Eq. 7)
2. To calculate the ligand binding free energy of the internal states, we need to utilize the all-atom coordinates. However, it is unreasonable to optimize the whole protein structure, as we intend to preserve the high-energy DOFs. Moreover, full optimization is impractical due to the unreasonable interatomic distance issue mentioned above. One approach is to freeze the protein conformation and place the ligand at a certain distance from the binding site. However, if a flexible model is desired, we suggest that users freeze (constrain) most part of the protein while keep the binding site (e.g., of the TS structure) and the ligand flexible (Figure 6), depending on your specific needs and available computational resources. Alternatively, you can also repair the unreasonable bond distances and angles using our raw structure as a template, thereby eliminating atomic collisions and obtaining a reasonable structure that allows reading and simulation. We provide demo in the standard package.
3. One could also perform other types of calculations, such as QM/MM, potential of mean force (PMF) calculation, EVB calculation, etc. However, one thing to keep in mind is that you should not push your protein out of the high energy state, or the database lost its meaning. We have to constrain certain DOFs that traditional sampling methods do not like, while making the sampling process feasible.

**Figure 5.**
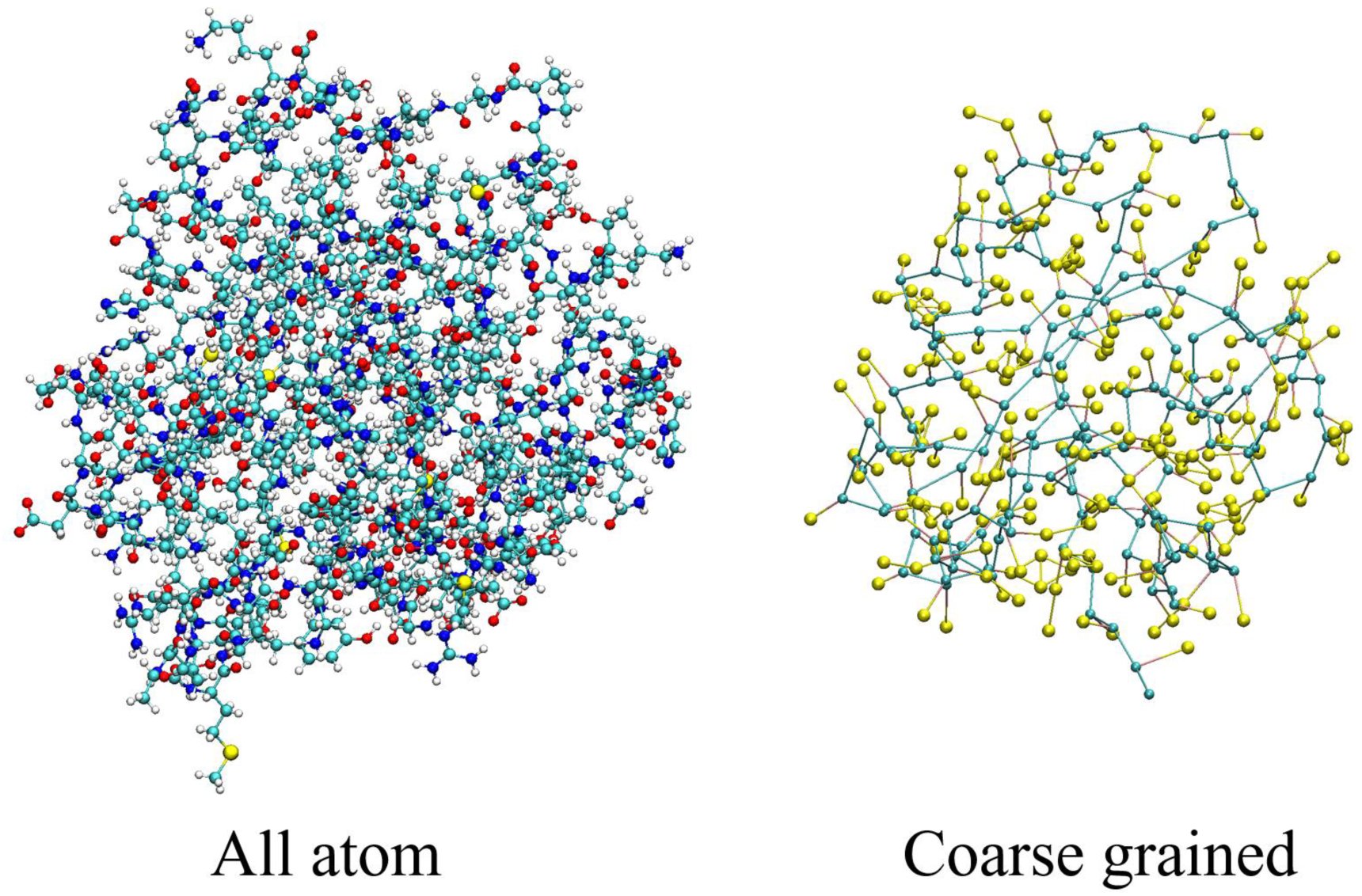
Visual comparison of the all-atom and coarse-grained representations. The protein is shown in all-atom CPK representation on the left and in CG representation on the right.

**Figure 6.**
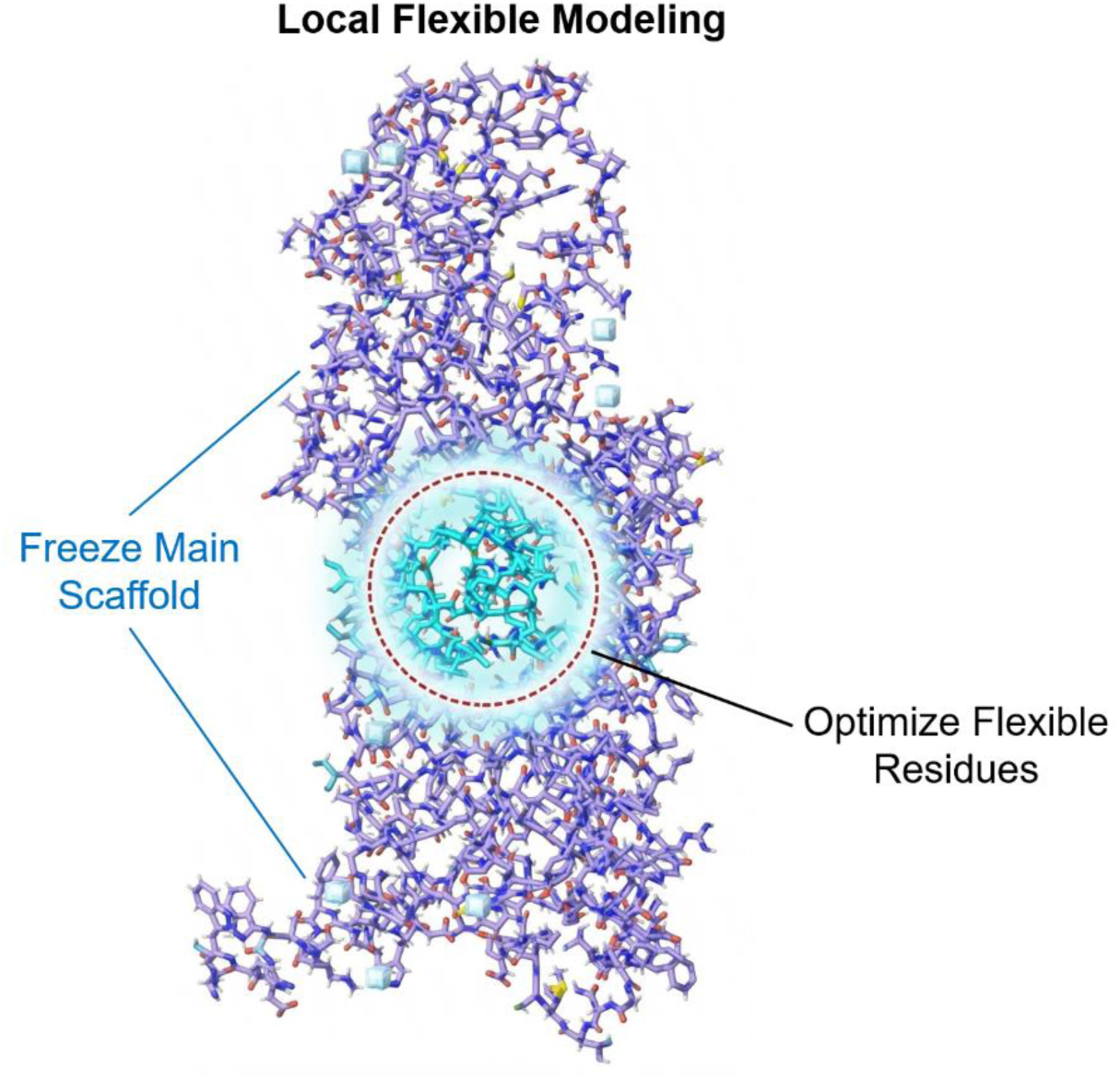
Local optimization restricted to the binding pocket region, while the rest of the structure is kept frozen or restraint.

### Examples of usage

#### Novel Pocket Pattern Identification

First of all, we show how this dataset can be used to identify new pockets that emerge and how old pockets varies during the activation process. As the protein undergoes conformational changes from the IAS toward the AS, transient pockets open up that are not present in the IAS. These pockets are novel, inaccessible to existing experimental techniques and unavailable in other datasets. By leveraging the full trajectory of the activation process, our dataset enables the discovery of such novel pocket patterns, offering new opportunities for targeting previously undruggable proteins. A representative example is illustrated in Figure 4. Pocket changes of all proteins in PAM-DB version 1.0 are shown in Figure 7 For complete movies, check website https://www.momedpamdb.com/en.

**Figure 7.**
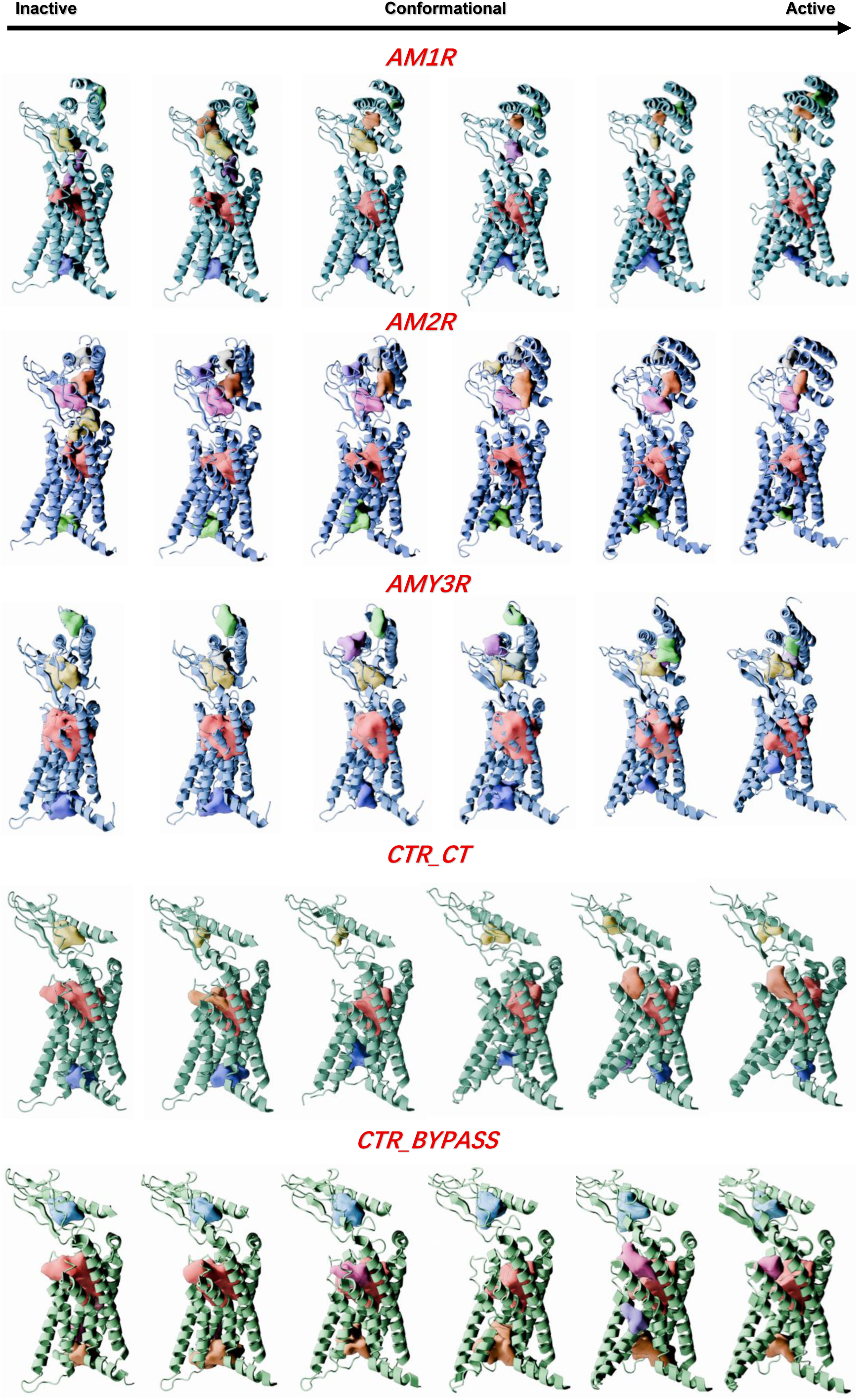
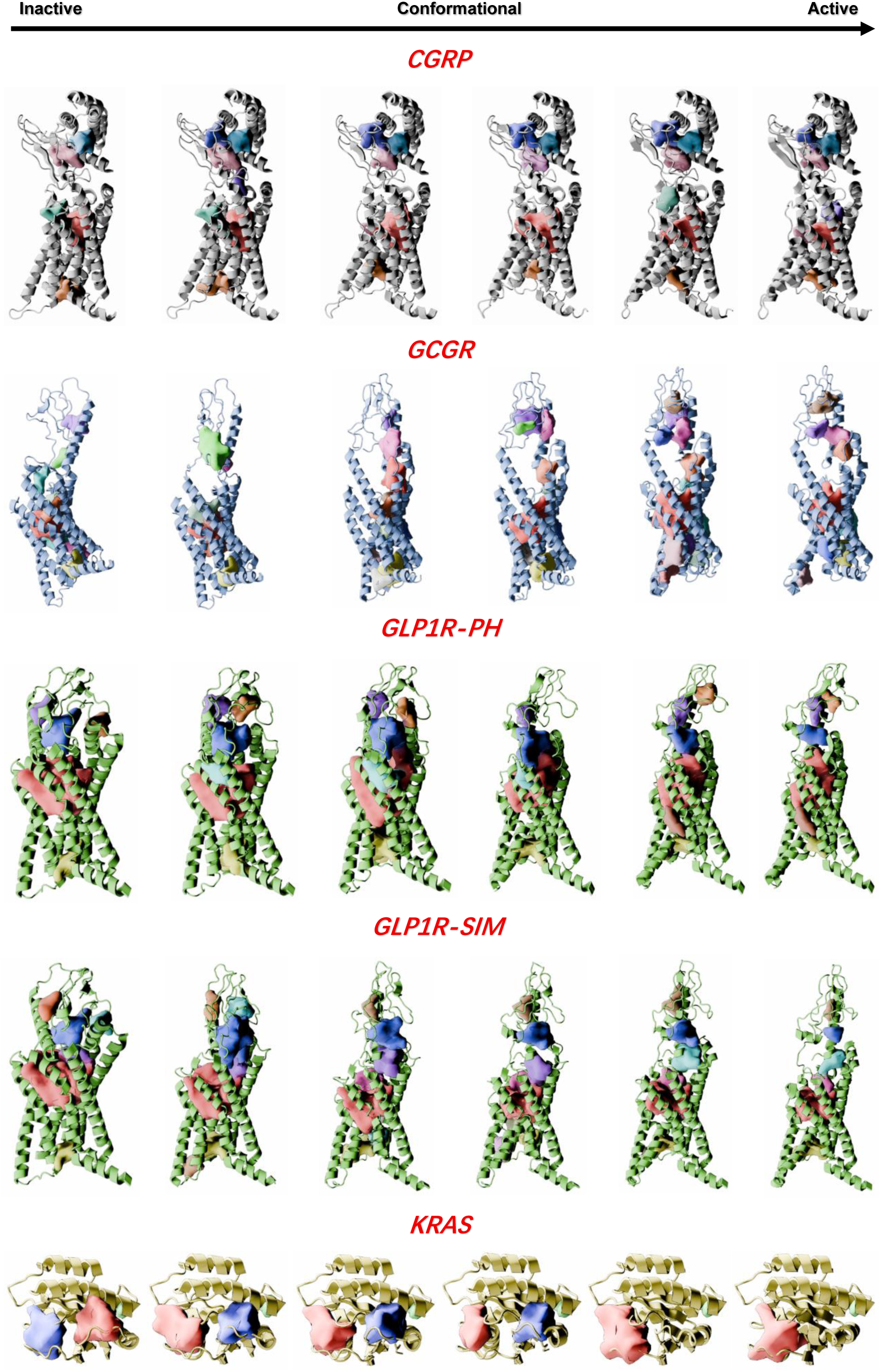
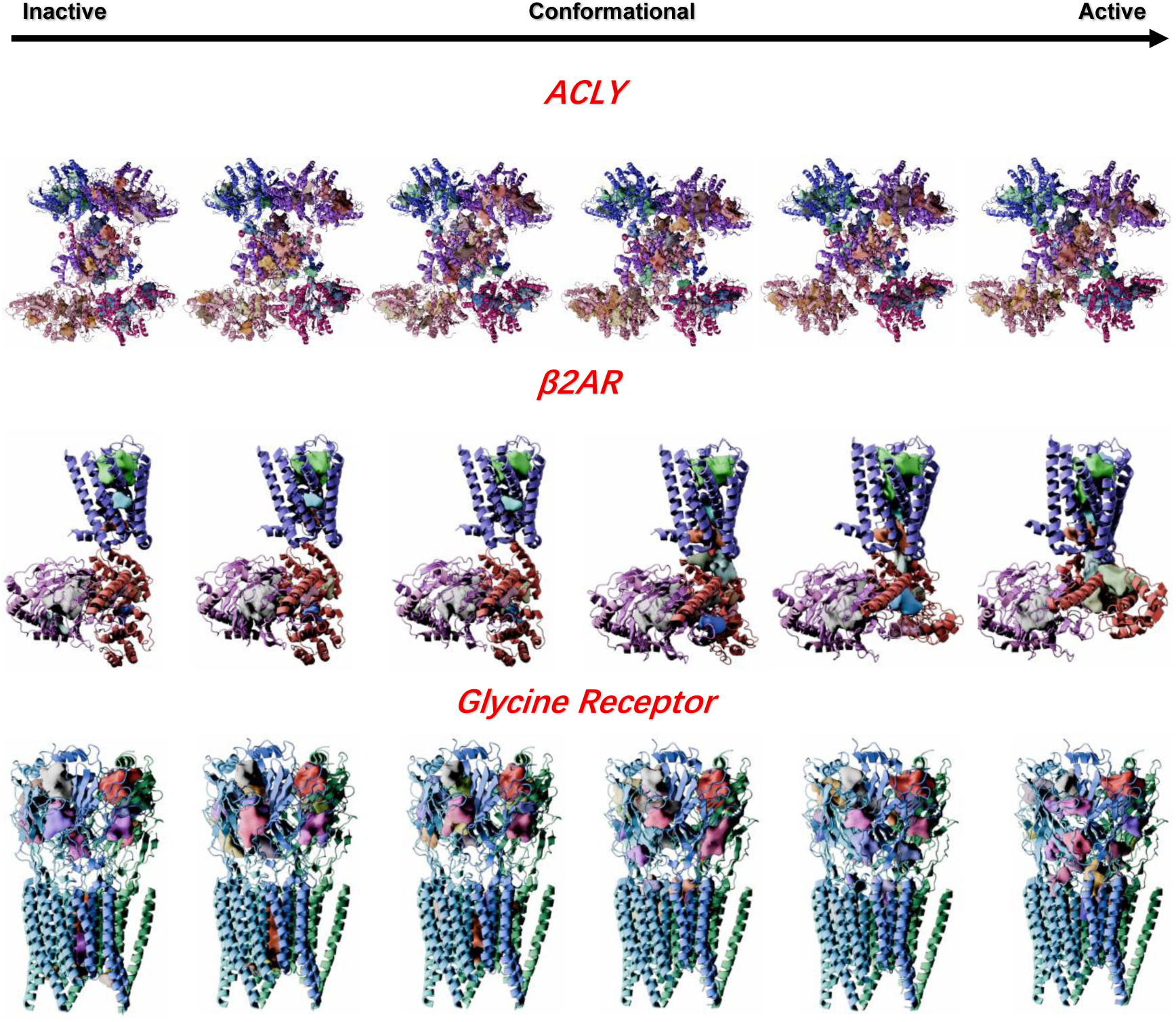
Overview of all pocket changes in PAM-DB 1.0. Proteins are depicted in cartoon representation, while pockets are rendered as surfaces. For complete movies, visit https://www.momedpamdb.com/en.

#### Biased Antagonist Design for An Undruggable Target

Then we present an example of how we used this workflow to design a Best-in-Class (BIC) partial inhibitor candidate for the “undruggable” target GCGR with novel skeleton.

GCGR is a prototypical Class B GPCR that is activated by glucagon and subsequently interacts with the stimulatory G protein. This interaction stimulates adenylate cyclase, leading to an increase in intracellular cyclic adenosine monophosphate (cAMP) levels.^13^ The overall activation mechanism of GCGR have been well defined as shown in Figure 8.

**Figure 8.**
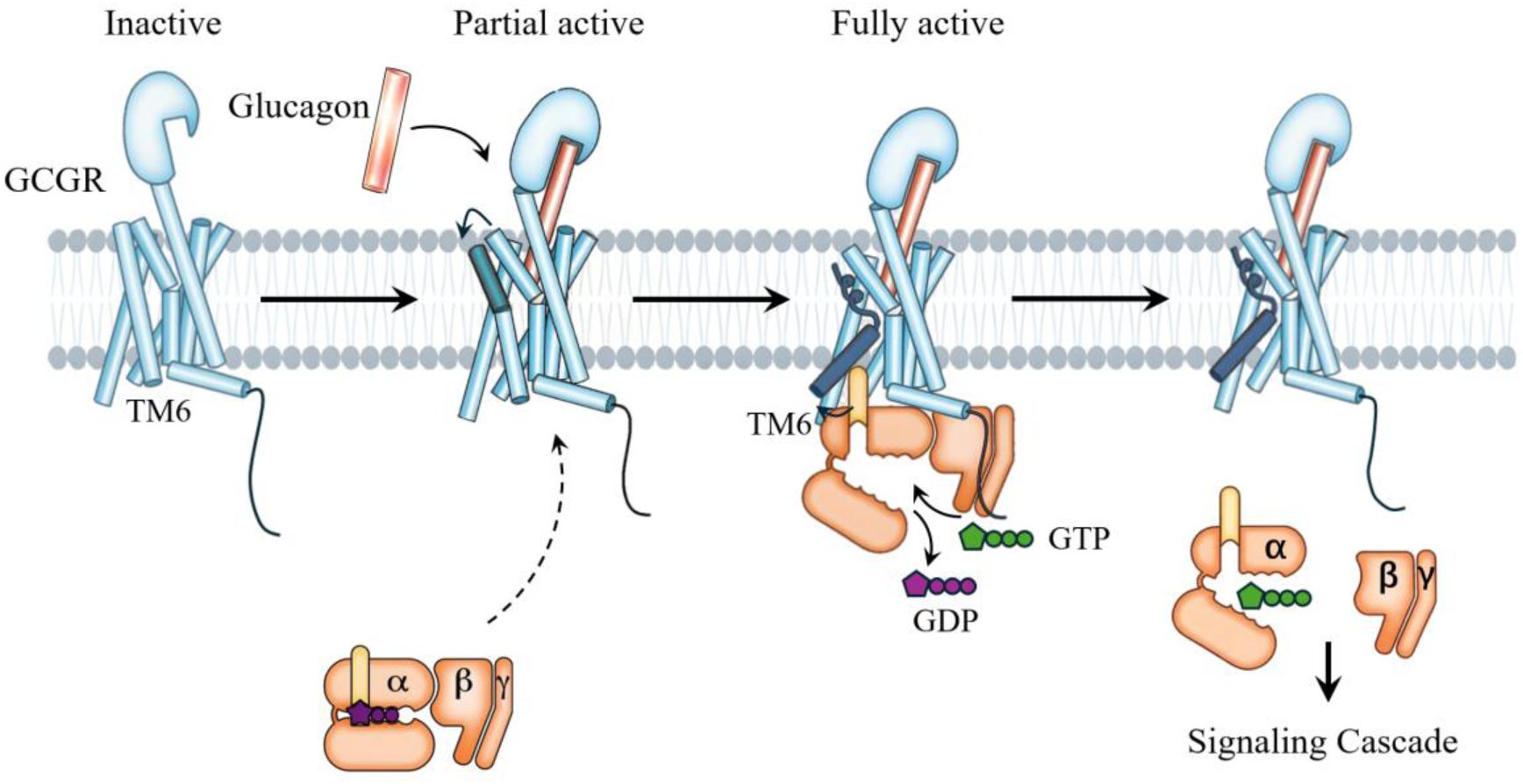
Schematic description of the GCGR activation pathway.

Despite numerous patents and extensive research efforts on GCGR, no successful inhibitor drug has reached the market to date (agonist already exist). Three clinical candidates have entered trials, but they share considerable structural similarities, reflecting a lack of molecular novelty (Figure 9). Moreover, these compounds have exhibited significant side effects, including elevated cholesterol levels. Our analysis suggests that the failure of these candidates is primarily rooted in toxicity issues arising from the dual signaling pathways activated by GCGR.

**Figure 9.**
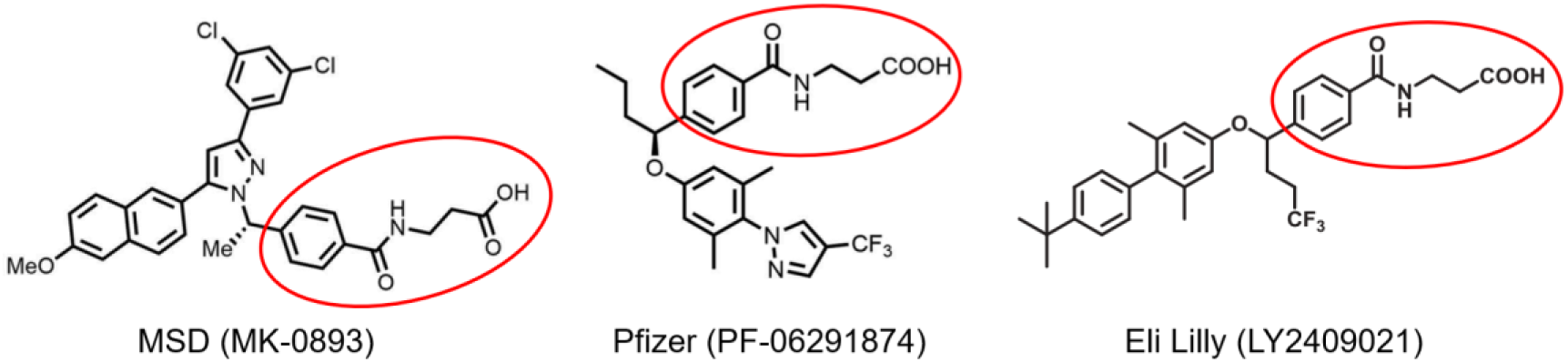
Structural comparison of three GCGR molecules that have entered clinical trials. The regions highlighted in red indicate conserved structural motifs, demonstrating their high structural similarity.

Upon activation, GCGR triggers two independent signaling cascades (Figure 10A): the canonical cAMP-PKA pathway via Gαs, which drives hepatic glucose output, and the β-Arrestin-dependent pathway, which mediates receptor desensitization, internalization, and ERK/MAPK signaling. The β-Arrestin pathway plays a protective role in maintaining liver metabolic homeostasis—its disruption leads to increased VLDL secretion, impaired hepatocyte regeneration, and rebound hyperglycemia. Traditional pan-GCGR inhibitors that block both pathways simultaneously achieve potent glucose-lowering effects but disrupt this delicate balance, explaining the toxicity observed in clinical candidates. Therefore, next-generation therapeutics must adopt a biased design strategy (Figures 10B and Figure 10C). Rather than fully inhibiting GCGR, ideal drug candidates should selectively block the cAMP-PKA pathway while preserving β-Arrestin function, thereby achieving efficacy without compromising metabolic safety. This requires in-depth mechanistic studies of how the two different signaling pathways are activated, including their conformational changes, energy variations, and other kinetic properties during the activation process, precisely the kind of information our dataset aims to provide.

**Figure 10.**
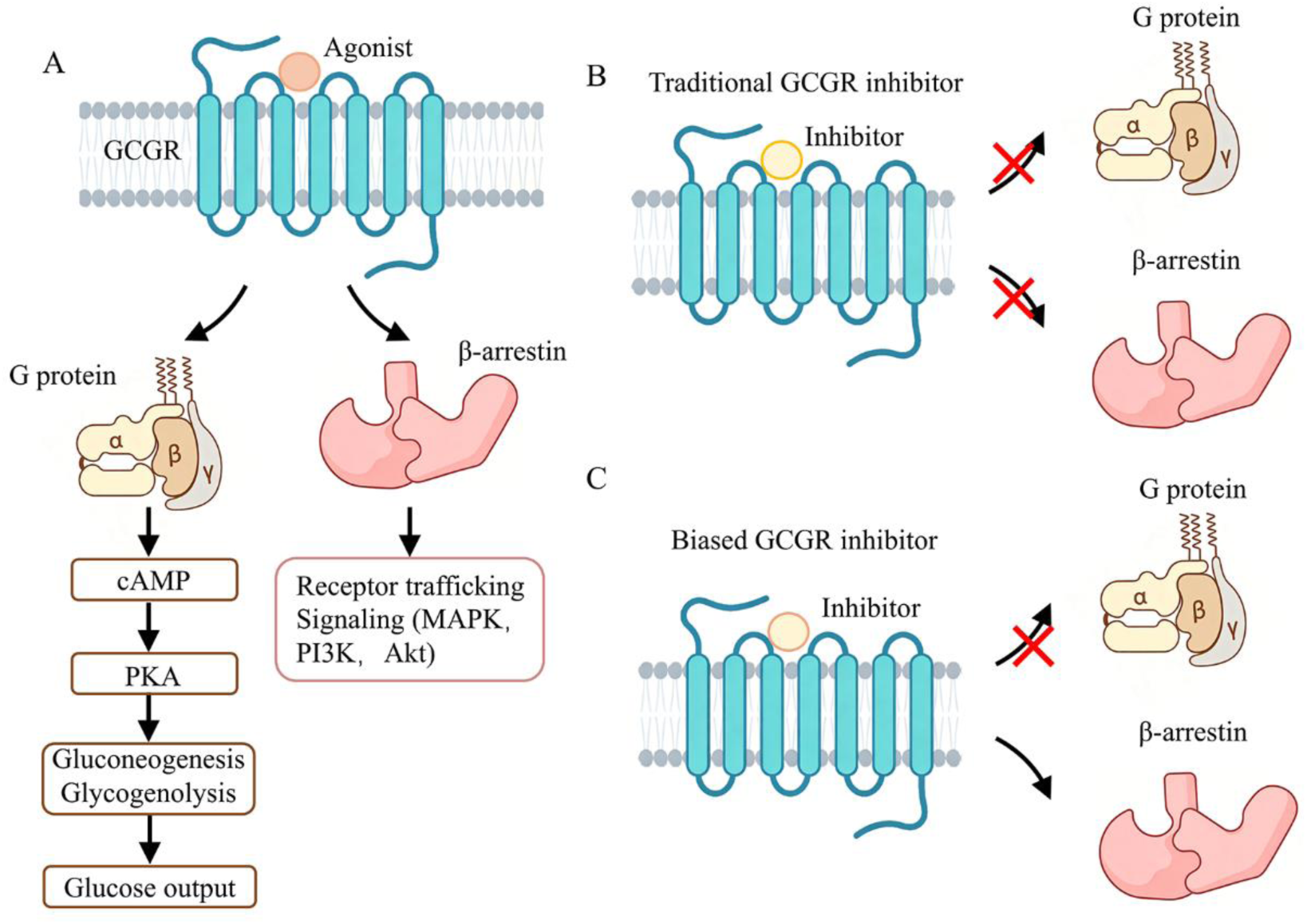
(A) Canonical GCGR signaling pathways via G protein and β-arrestin. (B) Traditional inhibitor blocks both G protein and β-arrestin pathways. (C) Biased inhibitor selectively inhibits G protein-mediated signaling while maintaining β-arrestin recruitment.

The overall workflow is illustrated in Figure 11. First, we systematically investigated the activation process of the GCGR protein, covering the conformational changes from the IAS to the AS, the coupling of G protein binding with GDP release, and the complete process of β-arrestin binding. Through computational analysis, we obtained key kinetic information—such as energy barriers and transition state conformations— along these processes, and identified several residues critical to the activation process via energy decomposition analysis.

**Figure 11.**
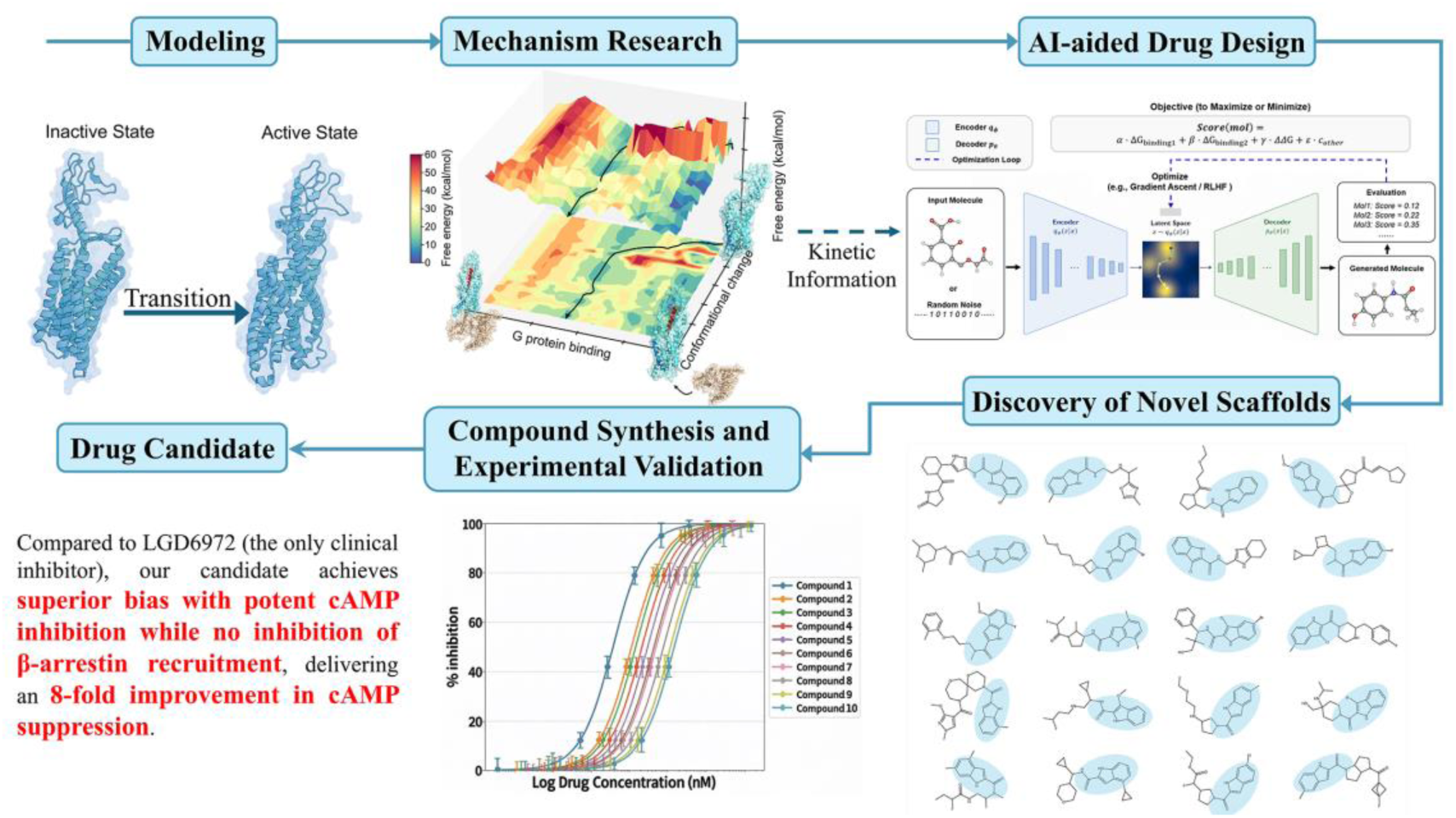
Overall workflow for the design of biased inhibitors targeting GCGR. The figures shown are schematic representations only and do not represent final experimental data, as they involve matters related to pending patent applications and future publications.

Then, based on these mechanistic insights, we adopted a data-driven iterative training strategy. Specifically, using large molecular databases such as REAL and ZINC as the initial chemical space, we evaluated the reaction energy barriers of these molecules, constructed a training dataset containing molecular structures and their corresponding barrier information, and trained an initial molecular generation model accordingly. The model was then used to generate new candidate molecules, whose effects on the target energy barrier were further evaluated. The newly obtained “molecule structure–energy barrier” data were continuously fed back into the training process to iteratively update the model, expanding the training data and improving model performance.

Regarding the optimization objectives, we set elevating the target energy barrier as the core optimization goal (i.e., inhibiting GCGR activation), while incorporating drug-likeness (QED), molecular diversity, logP, and synthetic accessibility into the reward function to achieve multi-objective optimization of the generated molecules. After candidate molecule generation, post-processing and barrier evaluation were performed to select the top 100 molecules with the greatest ability to elevate the target energy barrier for structural analysis. The results revealed several common structural features among these molecules, providing important references for subsequent molecular optimization.

On this basis, we further conducted post-processing on these candidate molecules, including computational evaluations such as molecular docking and MD simulations, and ultimately selected a batch of molecules for synthesis and activity testing. Experimental results identified several compounds with promising activity, and biased signaling assays were performed on these active molecules, yielding satisfactory results. Currently, our candidate molecules with novel backbone exhibit approximately 8-fold improved biased signaling selectivity compared to the only remaining clinical candidate (LGD6972). The designing process took three months and we synthesized fewer than 50 molecules.

#### Analysis of Existing Molecular Interaction Modes and Lead Optimization

Other than designing novel skeleton, another common practice in pharmacy companies is to modify and optimize existing drug candidates or molecules. Here we present an example using DPP1: based on mechanistic studies of existing inhibitors, we identified key residues that elevate the energy barrier and designed new interactions against these residues to optimize the compounds.

DPP1 (also known as CatC) is a cysteine protease that plays a central role in the maturation and activation of neutrophil serine proteases (NSPs), making it an important therapeutic target for neutrophil-mediated inflammatory diseases such as bronchiectasis. Currently, two representative inhibitors have entered clinical development: one is the approved drug Brensocatib, which forms a reversible thioimidate complex with Cys234 via its cyano group, effectively addressing the aortic toxicity caused by non-specific binding to aortic wall proteins observed with first-generation inhibitors; the other is BI 1291583, currently in Phase III trials, which exhibits preferential bone marrow distribution with significantly higher target-tissue exposure than prior DPP1 inhibitors, demonstrating superior tissue selectivity.

To further understand the inhibitory mechanisms underlying these clinical candidates, we first conducted mechanistic studies on DPP1, elucidating its complete conformational transition pathway from the IAS to the AS and identifying the key energy barriers and TS conformations along this process. Subsequently, we validated the mechanisms of the two inhibitors described above, confirming that both significantly elevate the activation energy barrier of DPP1. Building on this, we calculated the contribution of each residue to the overall energy barrier and identified those with the highest energy contributions. We then converted these key residues and their preferred interaction types into explicit interaction prompts, which were fed into the MechGen molecule generation module.^14^ Leveraging the model’s equivariant cross-attention mechanism, MechGen is able to precisely construct new pharmacophoric interactions at target residue sites through directed denoising and reshaping of local conformations, while preserving the geometric features of the molecular core scaffold, thereby enabling highly controllable lead compound optimization. Finally, experimental results demonstrated that our designed compounds exhibit excellent enzymatic activity and cellular potency, along with favorable metabolic stability and Best-in-class permeability. Notably, our lead compounds show a pronounced preference for bone marrow distribution, representing a marked improvement over the positive control BI 1291583—a molecule already known for its preferential bone marrow targeting. The designing process took two months and we synthesized fewer than 30 molecules. Due to commercial confidentiality, detailed information cannot be disclosed in this document.

## Conclusion

In summary, the kinetic determinants of protein function—including transition-state (TS) geometries, intermediate (IS) ensembles, and the associated free-energy barriers—remain the missing dimension in conventional structure-based drug discovery. Here we establish the first practical framework that systematically integrates these kinetic parameters into the rational design pipeline. Our database reveals that substantial conformational rearrangements along the activation trajectory give rise to transient pocket architectures that are entirely absent in both inactive and active end-states, thereby offering unprecedented opportunities for targeting previously “undruggable” proteins and discovering novel allosteric sites. More importantly, this approach transforms drug discovery from a static affinity-centric paradigm to a kinetic modulation strategy: it enables the design of candidates that not only bind tightly to the desired state but also selectively reshape the activation energy landscape to achieve a predetermined pharmacological outcome. We acknowledge that this work represents an initial step rather than a definitive solution. However, to our knowledge, it is the first practical realization of Activation Mechanism-Based Drug Design (AMBDD) — a paradigm that moves beyond structural snapshots to embrace the full dynamic choreography of protein function. We anticipate that this database will serve as both a discovery platform and a conceptual catalyst for the next generation of rational drug design.

- The v1.0 dataset includes: GLP-1R, GCGR, ACLY, GlyR, β2AR, KRAS, AM1R, AM2R, AM3R, CGRP, CTR.
- In following version 2.0, we intend to add additional protein, such as SHP2, M4R, etc. We welcome readers to share their thoughts.
- To see database availability and complete video of protein conversion in dataset, check https://www.momedpamdb.com/en.
- For commercial affairs contact
- For academic inquires contact

## Methods

All the data from this dataset is generated using open source software and homemade code.

## Reference

1. Ferrari, Á. J.; Dixit, S. M.; Thibeault, J.; Garcia, M.; Houliston, S.; Ludwig, R. W.; Notin, P.; Phoumyvong, C. M.; Martell, C. M.; Jung, M. D., Large-scale discovery, analysis and design of protein energy landscapes. Nature 2026, 1–11.

2. Bai, C.; Wang, J.; Mondal, D.; Du, Y.; Ye, R. D.; Warshel, A., Exploring the activation process of the β2AR-Gs complex. Journal of the American Chemical Society 2021, 143 (29), 11044–11051.

3. Bai, C.; Asadi, M.; Warshel, A., The catalytic dwell in ATPases is not crucial for movement against applied torque. Nature Chemistry 2020, 12 (12), 1187–1192.

4. Zhu, X.; Luo, M.; An, K.; Shi, D.; Hou, T.; Warshel, A.; Bai, C., Exploring the activation mechanism of metabotropic glutamate receptor 2. Proceedings of the National Academy of Sciences 2024, 121 (21), e2401079121.

5. Zhang, Y.; Zheng, Q.; Warshel, A.; Bai, C., Key Interaction Changes Determine the Activation Process of Human Parathyroid Hormone Type 1 Receptor. Journal of the American Chemical Society 2025, 147 (4), 3539–3552.

6. Zhang, Y.; Wu, K.; Li, Y.; Wu, S.; Warshel, A.; Bai, C., Predicting Mutational Effects on Ca2+-Activated Chloride Conduction of TMEM16A Based on a Simulation Study. Journal of the American Chemical Society 2024, 146 (7), 4665–4679.

7. Yan, J.; Chen, L.; Warshel, A.; Bai, C., Exploring the activation process of the glycine receptor. Journal of the American Chemical Society 2024, 146 (38), 26297–26312.

8. Bai, C.; Wang, J.; Chen, G.; Zhang, H.; An, K.; Xu, P.; Du, Y.; Ye, R. D.; Saha, A.; Zhang, A., Predicting mutational effects on receptor binding of the spike protein of SARS-CoV-2 variants. Journal of the American Chemical Society 2021, 143 (42), 17646–17654.

9. Liu, S.; Chen, H.; Zhu, X.; Ye, F.; Zhao, Y.; Qin, J.; Zheng, Y.; Wang, X.; Zhang, L.; Chen, H., Structural insights into the progressive recovery of α7 nicotinic acetylcholine receptor from nicotine-induced desensitization. Science Advances 2025, 11 (41), eadx4432.

10. Shi, D.; Zhu, X.; Zhang, H.; Yan, J.; Bai, C., Catalytic mechanism study of ATP-citrate lyase during citryl-CoA synthesis process. Iscience 2024, 27 (9).

11. Bai, C.; Warshel, A., Revisiting the protomotive vectorial motion of F0-ATPase. Proceedings of the National Academy of Sciences 2019, 116 (39), 19484–19489.

12. Lee, M.; Bai, C.; Feliks, M.; Alhadeff, R.; Warshel, A., On the control of the proton current in the voltage-gated proton channel Hv1. Proceedings of the National Academy of Sciences 2018, 115 (41), 10321–10326.

13. Mayo, K. E.; Miller, L. J.; Bataille, D.; Dalle, S.; Göke, B.; Thorens, B.; Drucker, D. J., International Union of Pharmacology. XXXV. The glucagon receptor family. Pharmacological reviews 2003, 55 (1), 167–194.

14. Du, H., W., Mingyang; Luo, Mengqi; Yang, Mengke; Yan, Yingchao; An, Ke; Zhu, Xiaohong; Warshel, Arieh; Hou, Tingjun; Bai, Chen, An Explicit Interaction-Prompted Diffusion Framework for High-Fidelity 3D Molecular Generation. Journal of the American Chemical Society 2026, Accepted.

